# Mechanical stress modulates source-to-sink partitioning and drought response in Arabidopsis

**DOI:** 10.64898/2026.03.04.709560

**Authors:** B. Lorena Raminger, Matías Capella, Facundo A Vanega, Raquel L. Chan, Julieta V. Cabello

## Abstract

Mechanical stimuli provoked by wind, soil compaction, rainfall, and biotic interactions strongly influence plant phenotype, growth, and development. Previous studies indicated that weight-treated *Arabidopsis thaliana* plants increase stem diameter, vascular bundle number, and seed yield, involving auxin, brassinosteroid, and strigolactone-related genes. In this work, we investigated how the mechanically induced increase in phloem area improves source-to-sink partitioning, while the increase in xylem area negatively affects long-term drought tolerance. Transcriptomic profiling confirmed a large-scale reprogramming of drought-responsive genes in treated plants. Moreover, quantification of sucrose and starch content highlighted an enhanced synthesis and carbohydrate transport, which ultimately and positively impacted lipid and protein contents in seeds. Using loss-of-function mutants, we demonstrate that the phloem loader SUC2 and exporters SWEET11, 12, and 16 are essential for the yield gains triggered by mechanical stress. Furthermore, mechanical treatment alters sugar metabolism. Overall, our findings indicate that weight treatment elicits a complex physiological response, in which sucrose transporters and starch metabolism play a crucial role in mediating its positive effects on seed quality and yield.

**Significance statement:** Mechanical cues are ubiquitous in natural environments, but their impact on plant carbon allocation and yield remains poorly understood. This study reveals that mechanical stress reshapes vascular architecture and carbohydrate transport, enhancing source-to-sink partitioning and seed quality in Arabidopsis. By identifying sucrose transporters and sugar metabolism as key mediators of mechanically induced yield gains, our findings provide mechanistic insight into how physical forces integrate with metabolic regulation influence plant productivity.

## INTRODUCTION

As sessile organisms, plants are unable to escape unfavorable environments and must adapt to diverse external challenges. These include abiotic stresses such as nutrient limitation, extreme temperatures, drought, and salinity, as well as biotic pressures caused by herbivory. Such factors can severely affect plant growth, development, and productivity, often resulting in significant yield losses (Khan et al., 2023; Kamatchi et al., 2024; Ramegowda et al., 2024). In addition to the abiotic and biotic stresses mentioned above, plants are also subjected throughout their life cycle to exogenous mechanical stress, defined as physical forces or pressures such as physical touch by passing animals, soil compaction, wind, raindrops, or competition with neighboring plants, all of which profoundly affect plant growth and morphogenesis (Brenya et al., 2022; Hansen et al., 2025). Some examples of mechanical stress are brushing, rubbing, touching, and changes in gravity (Brenya et al., 2022). The developmental and physiological responses triggered by these stressful cues are collectively called thigmomorphogenesis, an adaptive strategy that facilitates plants to cope with mechanical disturbances and holds significant relevance for agricultural practices (Jaffe and Biro 1976; Jaffe and Forbes, 1993). The thigmomorphogenic responses are mediated by multiple plant growth hormones, including auxin, ethylene, jasmonic acid, gibberelins, brassinosteroids, and strigolactones (Chehab et al., 2009; Brenya et al., 2022; Carrió-Seguí et al., 2024; Raminger et al., 2025). Mechanical stress not only affects growth, development, morphogenesis, and even survival of plants, but also induces the formation of cross-adaptation (Li and Gong, 2011; Brenya et al., 2022). For example, brushing on tomato plants increases chilling tolerance (Keller and Steffen, 1995), mechanical stimulation by increasing rotational speed improves heat, chilling, salinity, and heavy metal tolerances in tobacco suspension cells (Li and Gong, 2008; 2011; 2013), mechanical increases salt stress tolerance in tomato and *Ammopiptanthus nanus* plants (Capiati et al., 2006; Liu et al., 2024) or rubbing the stem with paper or stroking the shoot with the hand can enhance the ability of bean and maize to recover from drought (Jaffe and Biro, 1979). Among abiotic stresses, drought is one of the most critical, which causes an enormous reduction in crop yield (Kim et al., 2019). Plant responses and adaptation include a variety of cellular and molecular regulatory mechanisms required for short-term responses to prevent water loss *via* transpiration from guard cells and for long-term adaptations to acquire stress resistance at the whole-plant level (Nakashima et al., 2014; Takahashi et al., 2018).

Weight treatment, a type of mechanical stress, imparted on Arabidopsis stems modifies architectural features, leading to yield improvement. Seed weight in mechanically treated Arabidopsis was almost twice that of control, untreated plants (Cabello and Chan, 2019). Weight treatment increases the main stem width and the number of vascular bundles (VB). Plant vascular tissues are crucial for providing physical support and transporting water, sugars, hormones, and other small signaling molecules throughout the plant (De Rybel et al., 2015). Particularly, stem vascular tissue has two essential functions: it establishes the apical-basal axis for material transport, and provides continuity with the veins of lateral organs.

Sugars are central to plant growth and development, serving as energy sources and carbon skeletons for biomass production (Patrick et al., 2013). In plants, sucrose is the primary carbohydrate transported over long distances through the phloem, moving symplastically from mesophyll cells to the phloem parenchyma and subsequently being loaded into the phloem via apoplastic pathways mediated by SWEET and SUC transporters (Slewinski and Braun, 2010; Lemoine et al., 2013). In Arabidopsis, AtSWEET11 and AtSWEET12 facilitate sucrose efflux from phloem parenchyma cells, while the companion cell–localized transporter AtSUC2 is essential for efficient phloem loading and long-distance transport (Stadler and Sauer, 1996; Chen et al., 2012). Vacuolar SWEET transporters, such as AtSWEET16 and AtSWEET17, further contribute to intracellular sugar homeostasis (Chardon et al., 2013; Klemens et al., 2013). Proper regulation of carbohydrate partitioning is a major determinant of seed yield, and genetic manipulation of SWEET and SUC transporters has been shown to enhance seed filling and productivity in Arabidopsis and crop species, including rice (Braun, 2012; Chen et al., 2015; Fei et al., 2021).

Transported carbohydrates can be metabolized to obtain energy or stored as starch reserves. The synthesis and degradation of starch are regulated processes involving many enzymes. Among them, SS1 (STARCH SYNTHASE 1) participates in amylopectin synthesis, SS4 (STARCH SYNTHASE 4) promotes starch granule initiation, and GBSS is a granule-bound starch synthase (Szydlowski et al., 2011; Abt et al., 2020). ADG1 and ADG2 encode the small and large subunits of ADP-glucose pyrophosphorylase, which catalyzes the rate-limiting step in starch biosynthesis. PGM (PHOSPHOGLUCOMUTASE) catalyzes the reversible conversion of glucose 6-phosphate to glucose 1-phosphate. At night, starch degradation begins with the phosphorylation of glucan chains by GWD (GLUCAN WATER DIKINASE) and PWD (PHOSPHOGLUCAN WATER DIKINASE) (Ritte et al., 2006). This phosphorylation is thought to disrupt the structure of the outer chains, facilitating hydration and subsequent hydrolysis by β-amylases (BAM) proteins (Fulton et al., 2008). In Arabidopsis guard cells, the major starch-degrading enzyme is BAM1. When the day starts, BAM1 rapidly mobilizes starch with the chloroplastic AMY3, an ENDOAMYLASE that hydrolyzes α-1,4 bonds within glycan chains (Horrer et al., 2016). *BAM3* encodes a catalytically inactive protein located in plastids, where it may play a role in regulating starch metabolism (Fulton et al., 2008; Li et al., 2009).

Weight treatment increases the main stem width and the number of vascular bundles (VBs), leading to increased yield. Arabidopsis auxin influx carriers are crucial to achieving VB development (Cabello and Chan, 2019). Additionally, brassinosteroid (BS) signaling genes, such as *BZR1* and *BES1*, as well as strigolactone (SL) synthesis and response genes, including *MAX4* and *BCR1*, and a BRC1 target peptide, CLE44, are important for VB increase after weight treatment (Raminger et al., 2025). We wondered which genes are crucial to obtain the yield increase after mechanical stress. In this work, we investigated the role and distribution of carbohydrates, as well as that of SUC and SWEET transporters in the mechanical stress response leading to improved seed yield in Arabidopsis.

## RESULTS

### Weight treatment boosts seed yield via carbohydrate accumulation only in well-watered conditions

We previously reported that weight-treated plants show wider stems and a significant increase in the number of vascular bundles (VBs) compared to untreated control plants (Cabello and Chan, 2019). To determine which specific tissues are enlarged, different tissue areas were quantified. Xylem, phloem, and procambium areas from treated plants increased twice compared to untreated controls (Figures 1a and 1b). As xylem transports water and phloem photosynthates, we wondered if the observed anatomical changes affected such transport.

**Figure 1.**
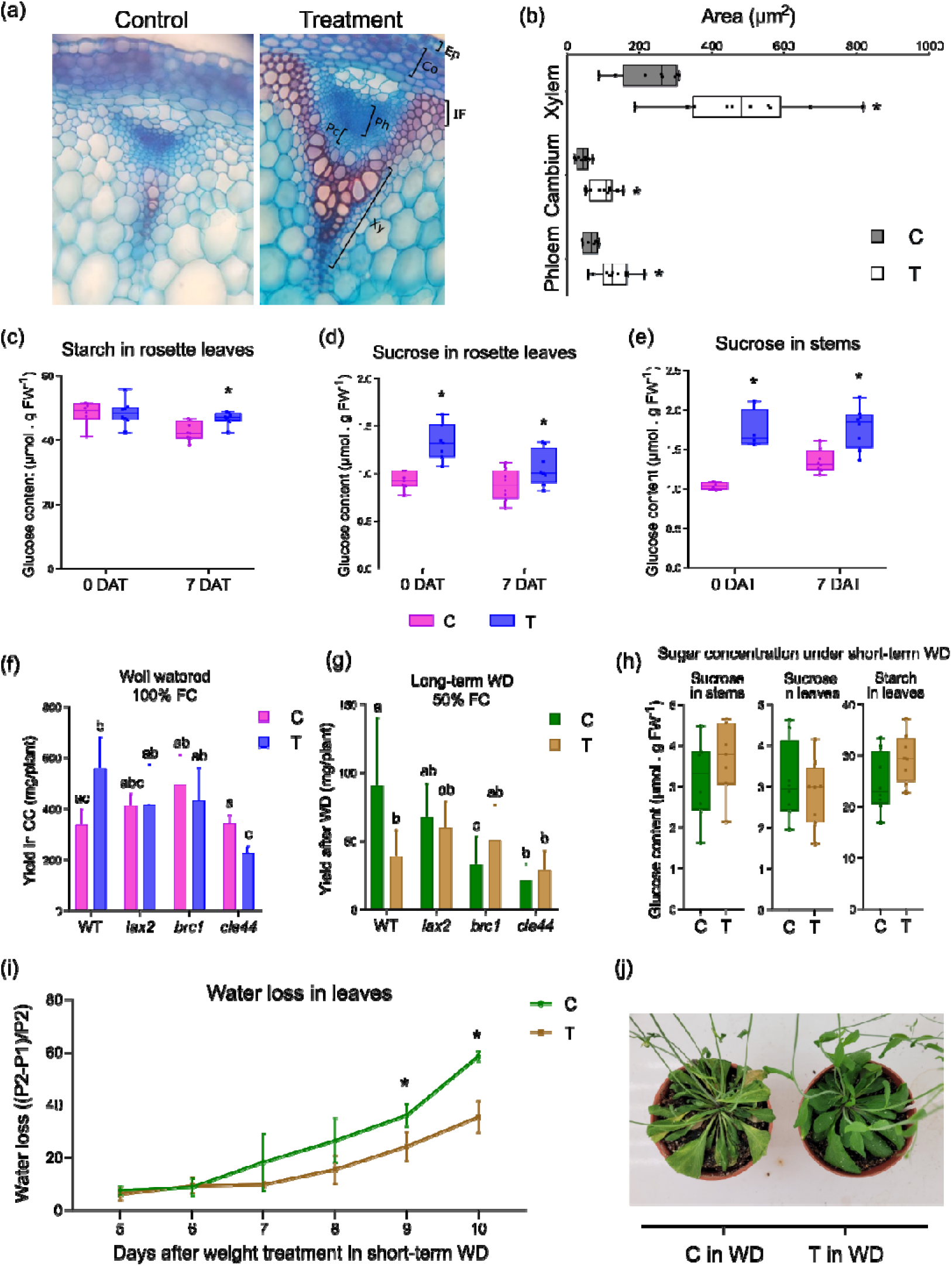
Weight treatment increases starch and sucrose accumulation, positively impacting seed yield in well-watered conditions but not under water deficit. **(a)** Illustrative picture of a vascular bundle of a stem cross-section stained with astra-safranin blue after 48 hours of weight treatment (right) and its control (left). (b) Quantitation of xylem, cambium, and phloem area (µm^2^) in stem cross-sections from control (C) and weight-treated plants (T) after 48 hours of treatment. (c) Starch content in rosette leaves, and (d) sucrose content in rosette leaves (e) and stem, immediately after weight removal (0 DAT) or after 7 days of weight removal (7 DAT) in control (C) and weight-treated plants (T). (f) Seed production of wild-type (WT) and *lax2*, *brc1*, and *cle44* mutant plants, in control (C) and weight-treated plants (T), under well-watered (WW) 100 % field capacity (FC) or (g) 50 % of FC (WD) conditions. (h) Sucrose content in stems and rosette leaves of weight-treated and untreated WT plants under short-term water deficit conditions. (i) Water loss evaluation during a short-term drought stress treatment applied to control (C) and weight-treated plants (T). (j) Illustrative photograph of C and T plants after nine days of short-term drought stress. Bars represent SEM (Standard Error of the Mean). Different letters indicate significant differences between means (P< 0.05, Two-way ANOVA with Tukey-Kramer post hoc test), and asterisks (*) indicate significant differences (P <0.05; Student’s t-test).

Considering the phloem function, we quantified sucrose and starch concentrations after 48 hours of weight treatment. Starch was quantified on the rosette (Figure 1c) and sucrose on the rosette and the stem (Figures 1d and 1e). Quantification was performed immediately after weight release (0 days after treatment, 0 DAT) and 7 (7 DAT) days after weight release on treated and untreated plants. At 0 DAT, sucrose concentration increased on the rosette leaves and in stems compared to untreated plants, whereas starch content remained unchanged (Figure 1c-e). At 7 DAT, rosette leaves had more starch and sucrose than controls, and the stems kept the sucrose increase in treated plants compared to controls (Figure 1c, 1d, and 1e).

Given that an increase in phloem area may enhance the concentration of transported sucrose, we evaluated whether an increase in xylem area could affect water transport and drought stress responses. We performed two different drought assays. Firstly, we evaluated the seed yield of plants subjected to long-term drought treatments, with field capacities (FC) maintained at 50%, 60%, and 80%. At the end of the treatment, weight-treated WT plants grown at 50% FC showed a significant reduction in seed yield compared with untreated plants, while mutant lines *lax2*, *brc1*, and *cle44*, that fail to increase stem diameter or vascular bundle number in response to mechanical stress, did not show changes in seed yield under 50% FC conditions (Figure 1g). These results indicate that an increased xylem area may negatively affect drought tolerance. At 60% FC, weight-treated WT plants displayed only a slight reduction in seed yield, whereas no yield differences were observed between treated and untreated WT plants at 80% FC (Figure 1f and Supplementary Figures 1a and 1b). During these assays, the plants remain viable. The other drought assay was designed to induce lethality and consisted of complete water withholding. Pots were saturated with water on the same day the weight treatment began. Leaf samples were collected during the drought treatment period to assess water loss and sucrose concentration. Weight-treated plants exhibited lower water loss than untreated plants after nine and ten days without irrigation (Figure 1i). Loss of turgor became evident in both groups during the final days of the treatment (Figure 1j). Despite the contrasting physiological responses, sucrose levels in stems and leaves were similar between weight-treated and control plants (Figure 1h). Overall, mechanical stress appears to enhance short-term drought tolerance while compromising long-term performance, as indicated by reduced seed production.

### Weight treatment induces changes in the expression of genes involved in drought tolerance

To assess the early transcriptional effects of mechanical loading, we analyzed the transcriptomic response of plants subjected to weight treatment at short time points. We performed a comparative RNA-seq analysis of Arabidopsis stems collected 6 hours after the onset of treatment, together with the corresponding control samples from untreated plants. In total, we identified 864 differentially expressed genes (DEGs; log_2_ fold change > 1.5 or < −1.5, false discovery rate < 0.05), including 556 upregulated and 308 downregulated genes, in weight-treated plants relative to untreated controls (Figures 2a and 2b; Supplementary Table S1). To identify the biological pathways affected by weight treatment, we conducted a gene ontology (GO) enrichment analysis of the DEGs. Among enriched GO terms, we found functional processes such as responses to insect and nitrogen compounds, aging, and responses to wounding and oxidative stress. Additional significantly enriched categories included genes involved in auxin and ethylene responses, as well as cell wall organization and biogenesis (Figure 2c; Supplementary Table S2). Consistent with our results (Figures 1i and 1j), GO term analysis revealed that thirty genes, most of them up-regulated, were associated with the response to water deprivation (Figures 2c and 2d), suggesting a strong correlation between mechanical and water deficit stress. Among regulated genes, *NAC019* was identified as a notable example among the DEGs related to water deprivation, as its overexpression has been shown to significantly increase drought tolerance (Tran et al., 2004). Accordingly, reverse transcription followed by quantitative PCR (RT–qPCR) confirmed that most of the selected drought-related genes were deregulated in weight-treated plants compared to controls (Figure 2e). Altogether, our transcriptomic analysis suggests that weight treatment modulates the expression of several drought-related genes, thereby enabling plants to respond more rapidly and to better tolerate severe short-term water deficit. We next asked whether different types of mechanical stimulation share common genetic networks underlying plant responses. By comparing the transcriptional profiles of plants subjected to weight (Ko et al., 2008; this work), touch (Xu et al., 2019), or hypergravity (Tamaoki et al., 2009), we found no significant overlap among either the upregulated or the downregulated gene sets (Supplementary Figure S2). These results suggest that each mechanical stress activates a distinct molecular response.

**Figure 2.**
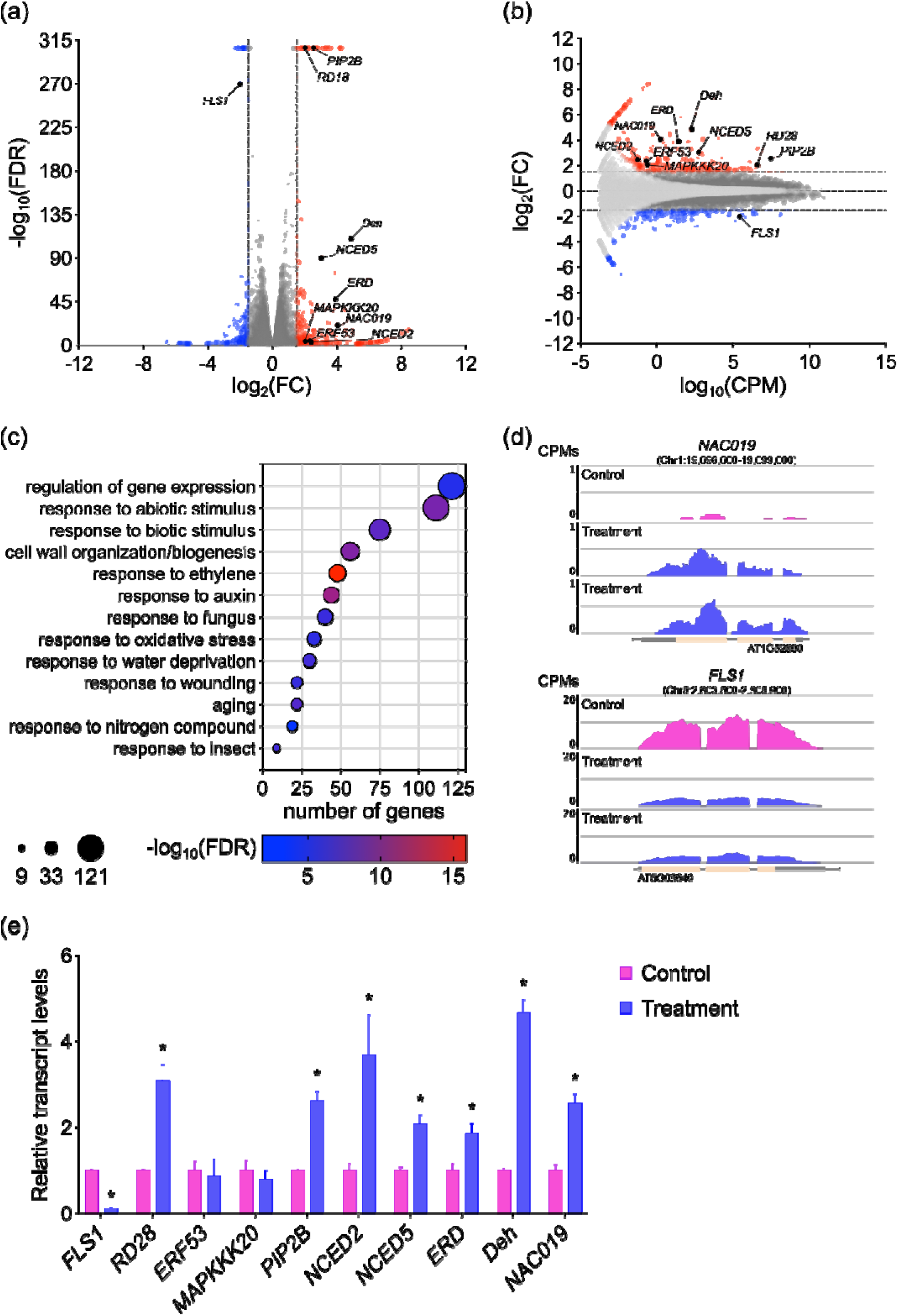
Weight treatment induces changes in gene expression and alters drought-responsive genes. (**a**) Volcano plot of RNA-Seq data from plants subjected to 6 hours of weight treatment versus untreated controls. Significantly upregulated (red) and downregulated (blue) genes are highlighted (log_2_(FC) > 1.5 or <-1.5 with FDR < 0.05, as determined using the edgeR framework). Selected transcripts of interest are labeled in black. (**b**) MA plot comparing treated plants with controls. The x-axis shows log_2_(CPM), and the y-axis shows log_2_(FC). All detected genes are shown, with those significantly upregulated or downregulated highlighted in red and blue, respectively. Genes displaying FDR > 0.05 are shown in light gray. (**c**) Functional enrichment analysis of genes that are significantly differentially expressed, as defined in panel (**a**). Point size reflects the number of genes in each category, and *p*-values are color-coded. (**d**) Coverage plots showing *RAP2-6* and *NAC019* transcript levels in untreated (pink) and treated (blue) plants. Reads are shown as counts per million (CPM), and genomic coordinates are indicated in base pairs (bp). (**e**) Relative transcript levels of different drought-responsive genes in untreated (pink) and treated (blue) plants, quantified by RT–qPCR. Data were normalized to *ACTIN2/8* transcript levels and are presented relative to untreated controls, whose value was set to one. All data represent mean ± SEM of *n* = 3 independent biological replicates. CPM, counts per million; FC, fold change; FDR, false discovery rate.

### Non-responsive genotypes fail to enhance sugar content and seed production

Weight treatment requires genes involved in auxin transport (*lax2*; Cabello and Chan, 2019) and in SL/BR pathways (*max4, brc1, bes1, bzr1, cle44*; Raminger et al., 2025), to enhance stem enlargement and seed yield. Mutants in those genes did not show stem thickening, increased vascular bundle number, or enhanced yield after treatment. To test whether this lack of yield increase was linked to sugar metabolism, we quantified starch in rosette leaves and sucrose in rosette leaves and first internodes in *lax2*, *brc1*, *cle44*, and WT plants after weight treatment.

Immediately after weight removal, starch content in rosette leaves did not differ between treated and untreated plants in any genotype. In contrast, sucrose concentration increased in the rosette leaves of weight-treated WT plants compared to untreated, whereas *cle44* mutants displayed higher sucrose concentrations than WT in untreated plants (Figure 3a). After seven days, both starch and sucrose concentrations increased in the rosette leaves of weight-treated WT plants compared to untreated, while none of the other genotypes showed differences in starch or sucrose concentrations (Figure 3a).

**Figure 3.**
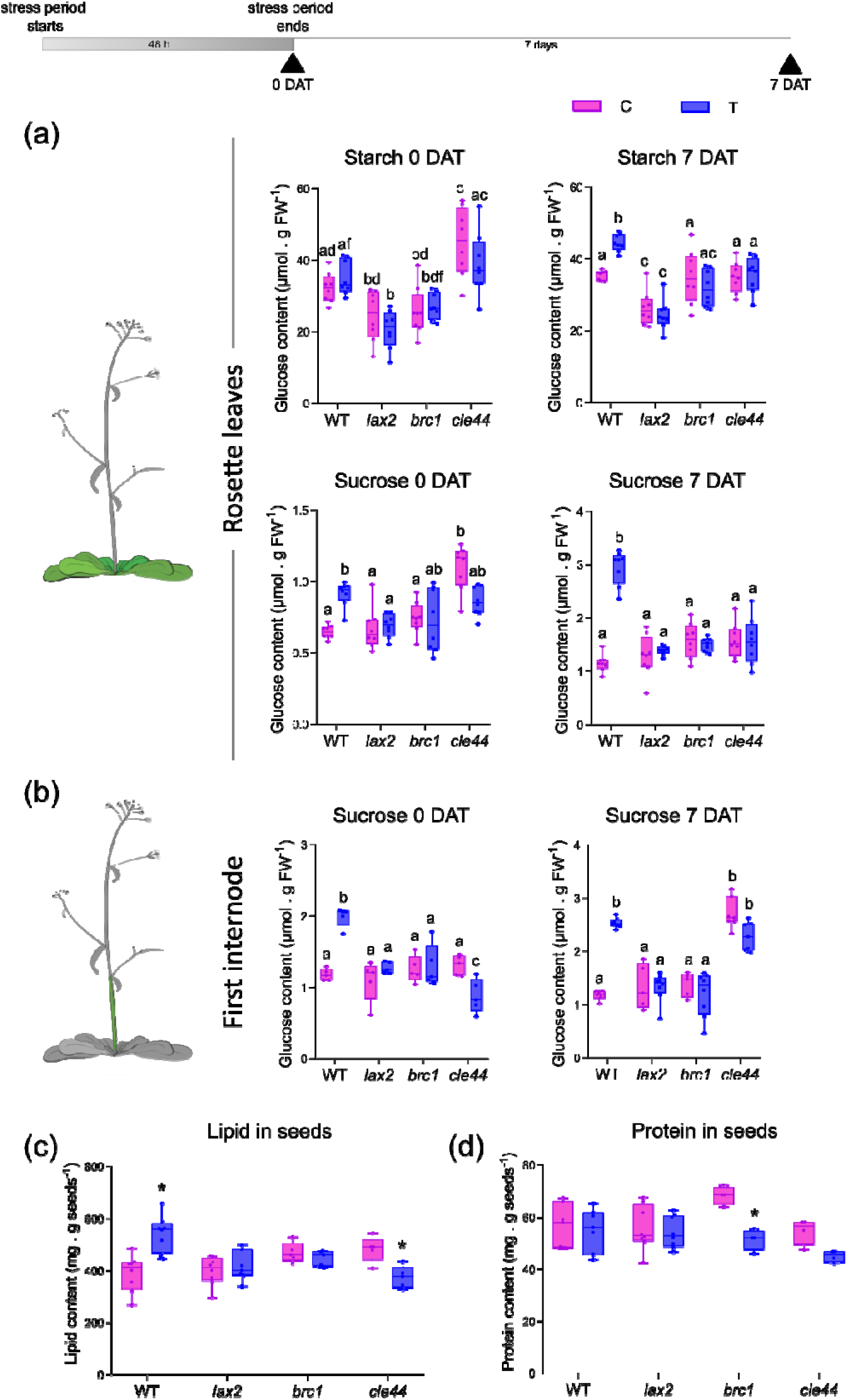
Carbohydrate metabolism is affected in weight-treated Arabidopsis plants but not in *lax2*, *brc1*, and *cle44* mutants. Starch and sucrose contents in (**a**) rosette leaves from untreated (C) and weight-treated (T) WT, *lax2*, *brc1,* and *cle44* mutant plants, immediately after weight removal (0 DAT) or after 7 days of weight removal (7 DAT). Sucrose content in (**b**) the first internode from untreated (C) and weight-treated (T) WT, *lax2*, *brc1,* and *cle44* mutant plants, immediately after weight removal (0 DAT) or after 7 days of weight removal (7 DAT). Lipid (**c**) and starch (**d**) content in seeds from untreated (C) and weight-treated (T) WT, *lax2*, *brc1,* and *cle44* mutant plants.

Sucrose concentration increased in the first internode of WT-weight-treated plants compared to untreated ones immediately after weight removal, whereas weight-treated *cle44* mutants exhibited less sucrose concentration in the first internode than in untreated *cle44*. Notably, sucrose concentration in weight-treated *cle44* plants was even lower than in untreated WT plants (Figure 3b). Seven days after treatment, sucrose concentration in the first internode remained higher in weight-treated WT plants compared to untreated ones, while none of the other genotypes displayed differences between treatments. Remarkably, untreated *cle44* plants accumulated more sucrose in the first internode than untreated WT plants (Figure 3b).

Based on the hypothesis that increased carbohydrate transport in weight-treated plants could influence seed quality, we analyzed lipid and protein contents in seeds. The results showed that lipid content was higher in weight-treated WT plants compared to controls (Figure 3c), supporting our hypothesis. In contrast, lipid content remained unchanged in *lax2* and *brc1* mutant plants, but was reduced in *cle44* plants, suggesting that these genes are required for weight-mediated increments in seed lipid accumulation and that CLE44 is essential for protection against treatment-induced lipid loss (Figure 3c). On the other hand, protein content in WT and *lax2* seeds was not affected by weight treatment. However, *brc1* mutants exhibited higher protein content under control conditions compared to WT, although their protein levels decreased following weight treatment. In *cle44* mutants, protein content also decreased after treatment (Figure 3d).

### AtSWEET and AtSUC transporters play crucial roles in carbohydrate transport during weight treatment

Given that sucrose concentration increased in internodes of weight-treated plants, we analyzed the expression of sugar transporters in rosette leaves and stems of weight-treated and untreated plants. Transcript levels of *AtSWEET9–15* and *AtSUC1–4* were assessed in rosette leaves, and *AtSWEET16–17* in the first internode after 48 hours of treatment. *AtSWEET12* and *AtSWEET16* were up-regulated in weight-treated plants compared to controls (Figure 4a and 4b).

**Figure 4.**
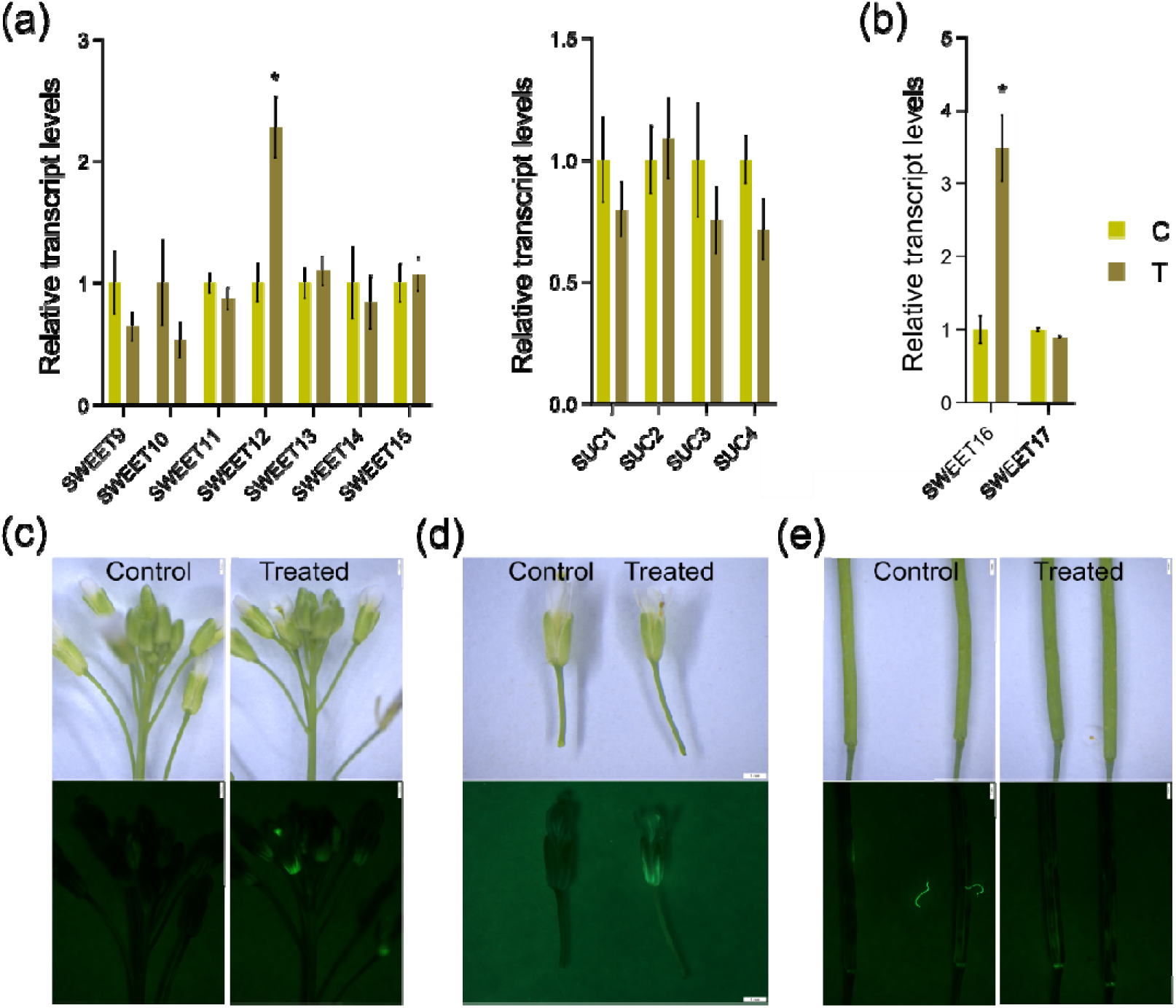
Sucrose transport is altered in weight-treated plants. (**a**) Transcript levels of *AtSWEET9, 10, 11*, *12*, *13*, *14*, *15*, (**b**) and *AtSUC1, 2, 3, 4* quantified by RT-qPCR in rosette leaves from control (C) and weight-treated (T) Arabidopsis plants. *AtSWEET16* and *17* quantified by RT-qPCR in the first internode from control (C) and weight-treated (T) Arabidopsis plants. The values were normalized with the one obtained in untreated plants (C). The experiment was performed with four biological replicates, obtained by pooling tissue from four individual plants and tested in triplicate. The bars represent SEM. Differences were considered significant and indicated with asterisks when the P-value was <0.05 (Student’s t-test). CFDA probe transported in (**c**) inflorescence, (**d**) mature flowers, and (**e**) siliques of weight-treated plants (T) compared to control untreated ones (C). Upper panels: bright-field images. Lower panels: CFDA probe transported in the same plants, visualized with a fluorescence microscope to detect the probe 5–10 min after CFDA application on the inflorescence base. Bars=1 mm

To assess whether weight treatment enhanced symplastic sucrose transport through the phloem, CFDA (a fluorescent dye) was applied to the base of cut inflorescences, mature flowers, and siliques of control and treated plants at flowering. Ten minutes after application, treated plants showed stronger fluorescence in flowers and siliques, indicating improved dye loading (Figures 4c–4e), indicating that weight-treated WT plants exhibit enhanced CFDA transport, likely reflecting the combined contribution of increased symplastic transport and the induction of SWEET transporters.

### SWEET 11, 12 and 16 are crucial to obtain seed increase after mechanical stress

To investigate the role of SUC and SWEET transporters in the mechanical stress response, *suc2*, *sweet11*, *sweet12*, *sweet11/12*, and *sweet16* mutants were subjected to weight treatment. Stem diameter increased approximately 50% (from 0.12 to 0.18 cm) across all genotypes after 48hours (Figure 5a). Similarly, the number of vascular bundles increased in all mutants following treatment (Figure 5b), indicating that these transporters are not required for the mechanical stress-induced changes in stem morphology. In contrast, seed production remained unchanged in treated mutants compared to untreated controls (Figure 5c). Together, these results highlight a critical role for *SUC2* and *SWEET11,12* y *16* in mediating seed yield enhancement under mechanical stress. Representative stem cross-sections confirmed the increase in vascular bundle number in all genotypes (Figure 5d).

**Figure 5.**
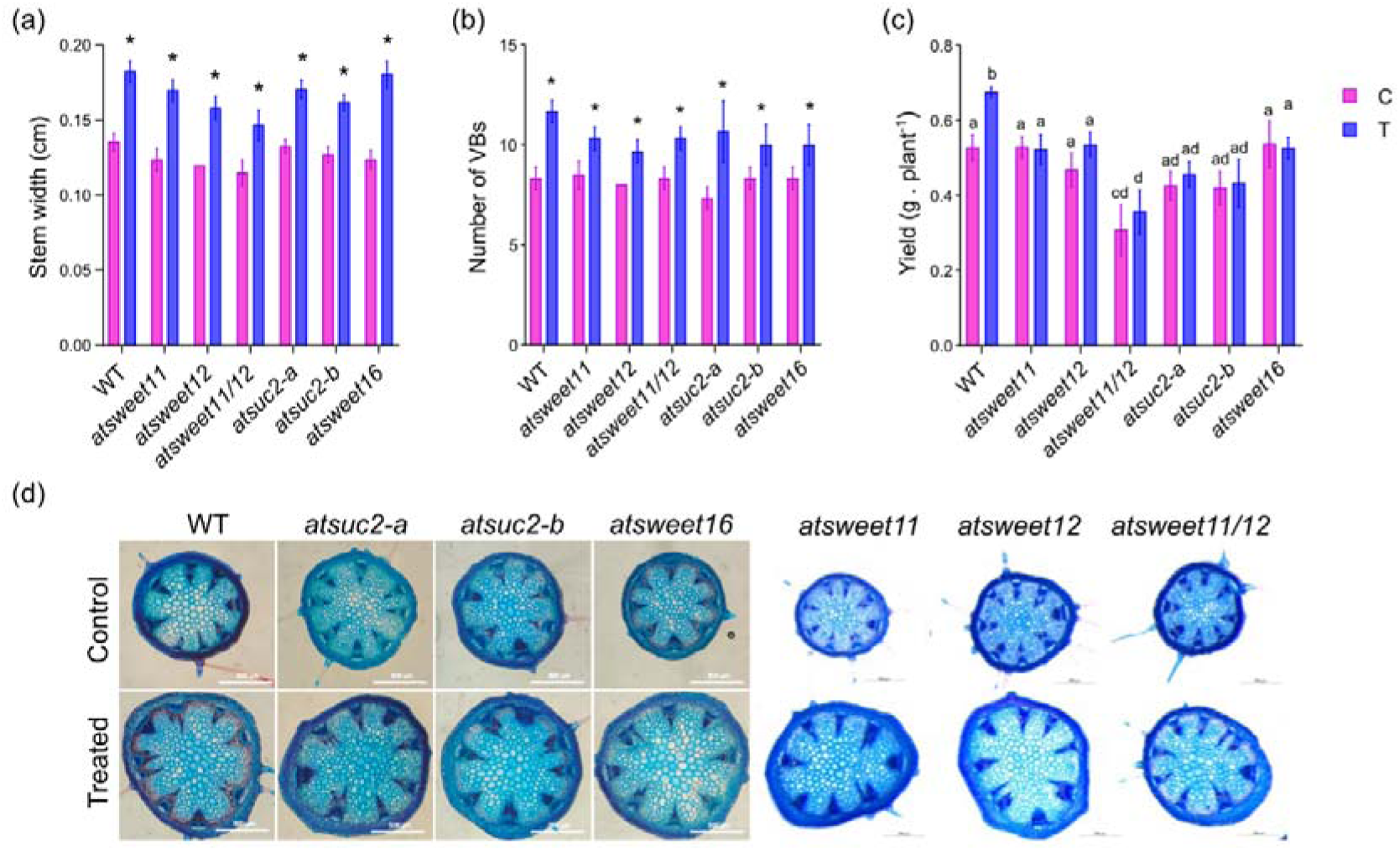
AtSWEET11, 12, 16, and AtSUC2 are necessary to achieve yield increase after weight treatment in Arabidopsis. First internode width (**a**) and number of vascular bundles (**b**) in the stem of *atsweet11*, *atsweet12*, *atsweet11/12*, *atsuc-2a*, *atsuc-2b,* and *atsweet16* mutant plants, and WT as control, measured 48 hours after weight treatment. Yield is measured as the weight of seeds (g) per plant under control conditions and after 48 hours of weight treatment. Cross-sections of the first internode of *brc*1 and *max4* mutant plants and WT as control after 48 hours of weight treatment (**d**). The bars represent SEM. Differences were considered significant and indicated with asterisks when the P-value was <0.05 (Student’s t-test).

### Starch metabolism genes are induced in weight-treated plants

Considering that weight-treated plants exhibited higher yield than controls (Cabello and Chan, 2019) and that sucrose levels were increased in stems and rosette leaves after weight treatment (Figures 1 and 3), we asked whether genes encoding enzymes involved in starch synthesis and degradation were differentially regulated in response to weight application.

Although RNA-seq analysis did not reveal significant changes in the expression of starch metabolism-related genes, we analyzed the transcript levels of key starch metabolism-related genes in rosette leaves of weight-treated and control plants at three different time points (Figure 6a), which differ from those used in the RNA-seq analysis (performed after 6hours of weight treatment). Weight treatment was initiated at 12:00 am after 6 hours of light. Samples were collected after 30 hours of treatment (matching with the end of the light period on the first day of treatment), 36 hours (matching with the onset of the night period), and 48hours (matching with the middle of the light period). After 30 hours, *PGM*, *ADG1*, *SS4*, and *GWD* were upregulated in treated plants compared to controls, indicating activation of both starch biosynthesis and degradation pathways (Figure 6b). After 36hours of weight treatment, at the onset of the night period, only *GWD* remained upregulated, suggesting that starch degradation was specifically induced in weight-treated plants (Figure 6c). Finally, after 48 hours of weight treatment, during the middle of the light period, *PGM*, *ADG1*, *GBSS*, and *SS4* were upregulated, consistent with activation of starch biosynthesis in these plants (Figure 6d). Together, these results indicate that weight treatment rapidly modulates the temporal regulation of starch metabolism in a light-dependent manner.

**Figure 6.**
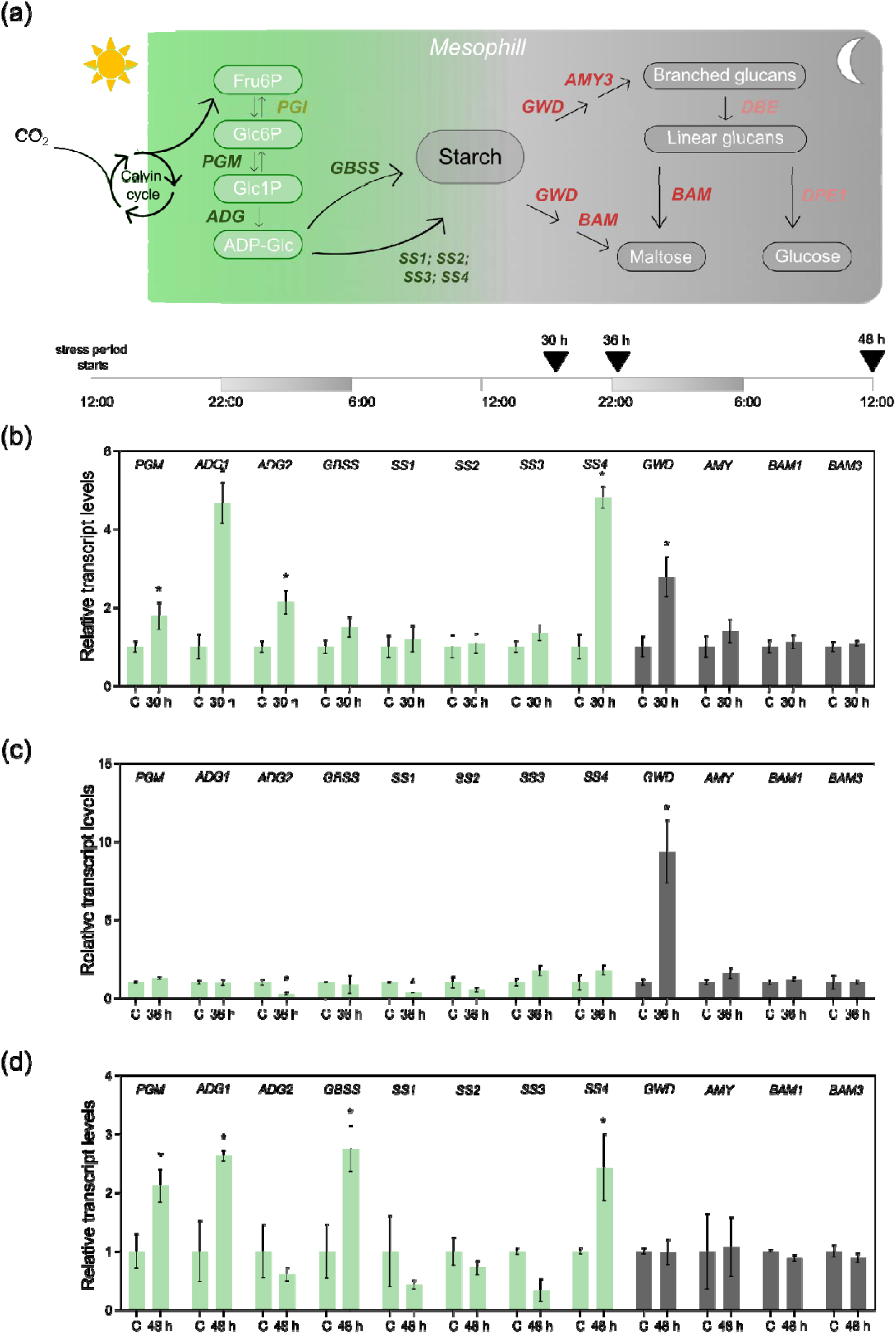
Transcript levels of genes involved in starch metabolism are differentially modulated in weight-treated plants compared to controls. (**a**) Illustrative scheme showing the functions of genes encoding enzymes involved in starch synthesis and degradation, along with a schematic representation of the stress assay performed on Arabidopsis plants. Grey bars indicate night periods and white bars indicate light periods. Black arrows mark the time points at which samples were collected (**b**) *PGM*, *ADG1*, *ADG2*, *GBSS*, *SS1*, *SS2*, *SS3*, *SS4*, *GWD*, *AMY3*, *BAM1,* and *BAM3* transcript levels quantified by RT-qPCR in weight-treated plants after 30 hours, (**c**) 36 hours, and (**d**) 48 hours. The values were normalized with the one obtained in WT. The experiment was performed with four biological replicates, obtained by pooling tissue from four individual plants and tested in triplicate. The bars represent SEM. Differences were considered significant and indicated with asterisks when the P-value was <0.05 (Student’s t-test).

## DISCUSSION

Plants often experience mechanical stress due to increased body weight and surrounding cell growth and expansion. Developmental plasticity enables them to modulate their phenotype in response to environmental conditions, thereby influencing biomass and seed yield efficiency (Sultan, 2010; Du and Jiao, 2020; Sampathkumar, 2020).

Mechanical stress has been shown to influence vascular development (Agustí and Blázquez, 2020). Weight application on the main stem of Arabidopsis not only increases the number of vascular bundles (Cabello and Chan, 2019) but also promotes vascular tissue expansion within each bundle (Figure 1). This represents the first experimental system directly linking mechanical stress-induced modifications in stem architecture and anatomy with a key agronomic trait, enhanced seed production. These observations are consistent with other systems in which mechanical stress drives architectural remodeling through the inherent developmental flexibility of plant tissues. For example, when mechanical forces are applied to mature leaves, the existing venation pattern can reorient in the direction of the applied stress (Bar-Sinai et al., 2016). Similarly, the application of weight to the main stem of Arabidopsis promotes secondary growth and vascular differentiation (Ko et al., 2004). Such mechanically induced architectural changes rely on the intrinsic plasticity of plant tissues, enabling continuous structural and developmental adjustments in response to external mechanical cues. This capacity provides an adaptive advantage by allowing plants to optimize transport efficiency and mechanical stability under fluctuating environmental conditions.

Mechanical stress affects multiple aspects of plant biology, including growth, development, morphogenesis, and survival, and may also promote cross-adaptation, enabling plants to better tolerate other abiotic stresses such as chilling, salinity, or drought. (Li and Gong, 2011, Brenya et al., 2022). In this work, weight-treated plants responded better to water-deficit conditions, as they lost less water under short-term drought treatment. The up-regulation of drought-responsive genes to prevent water loss *via* transpiration at early stages of mechanical treatment could explain the tolerance displayed by weight-treated Arabidopsis plants under this stress. In addition, the observed increase in xylem area under mechanical stress may contribute to this response. A larger xylem network enhances water transport efficiency and helps to maintain hydraulic conductivity under limited water availability (Sperry et al., 2007; Cornelis and Hazak, 2021). Thus, the mechanically induced expansion of xylem tissues may represent an adaptive response that not only supports higher seed yield, but also transient resilience to drought conditions (Cornelis and Hazak, 2021). The previously reported increase in lignin content in weight-treated WT stems (Cabello and Chan, 2019) may further enhance the ability of plants to sustain the high negative pressures required for long-distance water transport. A similar phenotype was described for the *nut1* mutant in maize, which exhibits impaired water movement due to reduced protoxylem thickness that secondarily compromises metaxylem cell wall integrity (Dong et al., 2020). Compared with vegetative stages, reproductive development is more sensitive to abiotic stress, particularly during the seed and fruit set phase around fertilization (Kakumanu et al., 2012; Ruan et al., 2012). This is manifested by the phenotypic difference in the severity of their responses; abiotic stress often reversibly inhibits leaf expansion but causes substantial abortion of flowers, young seeds, and fruitlets, and hence irreversible yield loss (Boyer et al., 2007; Muller et al., 2011). However, despite the short-term drought response, weight-treated plants ultimately lost their drought tolerance at long-term drought assays and even produce less seeds at 50% field capacity. A plausible explanation is their increased reproductive sink demand. Weight-treated plants produce more siliques, each containing more seeds (Cabello and Chan, 2019), and enhance the flux of photoassimilates toward developing reproductive structures. However, under prolonged drought, photosynthetic activity is substantially reduced, limiting the supply of assimilates required to sustain this enhanced sink demand (Kaur et al., 2021; Qiao et al., 2024). As a consequence, the imbalance between source capacity and sink strength could lead to seed abortion or incomplete seed filling, ultimately reducing yield under long-term water-deficit conditions.

Transcriptomic data indicate that the largest groups of differentially expressed genes correspond to insect-response pathways and nitrogen-related processes, with additional but smaller sets of genes associated with aging, wounding, water deprivation, and oxidative stress. Regarding hormonal pathways, several ethylene- and auxin-responsive genes were enriched, a pattern also observed in tomato plants subjected to a similar mechanical stimulation (Castro et al., 2025). When comparing our transcriptome with that reported by Ko et al. (2004), we found minimal overlap, restricted to a few genes involved in stem development in Arabidopsis. Similarly, little overlap was detected when comparing our dataset with the transcriptomes generated under hypergravity (Tamaoki et al., 2014) or touching stimuli (Xu et al., 2019), suggesting that different types of mechanical stress, and possibly the distinct phenological stages at which the treatments were applied, elicit largely divergent transcriptional signatures.

Seeds from siliques of weight-treated plants exhibited higher lipid content than untreated ones, demonstrating that better sugar transport has an impact not only on seed weight but also on seed nutritional characteristics. Furthermore, sucrose transporters like AtSUC2, AtSWEET12, and 16 were up-regulated in weight-treated plants compared to controls and it was demonstrated that seed filling in *Arabidopsis thaliana* depend on the three sucrose transporters SWEET11, 12, and 15 (Chen et al., 2015), and mutant plants carrying insertions in *AtSWEET11* and *12* are defective in phloem loading (Chen et al., 2012). In agreement, the yield of the double mutant *sweet11;12* was reduced. This reduction is due to a decrease in the seed weight of *sweet11;12* (Chen et al., 2015). In the same way, transgenic rice expressing *AtSUC2*, which loads sucrose into the phloem under the control of the *PHLOEM PROTEIN2* promoter (*pPP2*), showed an increase in grain yield (Wang et al., 2015). *AtSWEET17* was down-regulated in weight-treated plants compared to controls. It was reported that when *AtSWEET17* expression is reduced, either by induced or natural variation, fructose accumulates in leaves, suggesting an enhanced storage capacity (Chardon et al., 2013). Parallelly, weight treatment induced the expression of genes encoding enzymes involved in starch synthesis and degradation. Although starch metabolism genes did not show significant differential expression in our RNA-seq at 6 hours, the dynamic regulation, observed at later time points, suggests a systemic reconfiguration of carbon metabolism in response to mechanical stress. Consistent with this, transcriptome profiling in tomato subjected to mechanical treatment revealed significant changes in the expression of genes involved in central metabolism, including carbohydrate pathways, following mechanical stimuli (Castro et al., 2025). Similarly, mechanical damage has been reported to alter primary metabolite levels and resource allocation in other plant species, indicating that mechanical cues can reallocate carbon resources beyond immediate structural responses (Chen et al., 2021). These observations support the idea that mechanotransduction triggers metabolic shifts that may underlie the temporal modulation of starch synthesis and degradation observed in our system and contribute to broader adaptive responses.

## Conclusion

Weight treatment needs LAX2 to assure auxin flux to obtain more vascular bundle development (Cabello and Chan, 2019), BES1 and BZR1 to increase BRs signaling and xylem differentiation, and BRC1 and CLE44 to regulate procambium cell division to obtain more vascular bundles after weight treatment. To achieve yield improvement, SUC2, AtSWEET11,12 and 16, and genes involved in starch synthesis and degradation pathways are crucial. Altogether, weight treatment modifies the expression of genes that belong to different pathways and are needed to reach the complex phenotype described (Figure 7).

**Figure 7.**
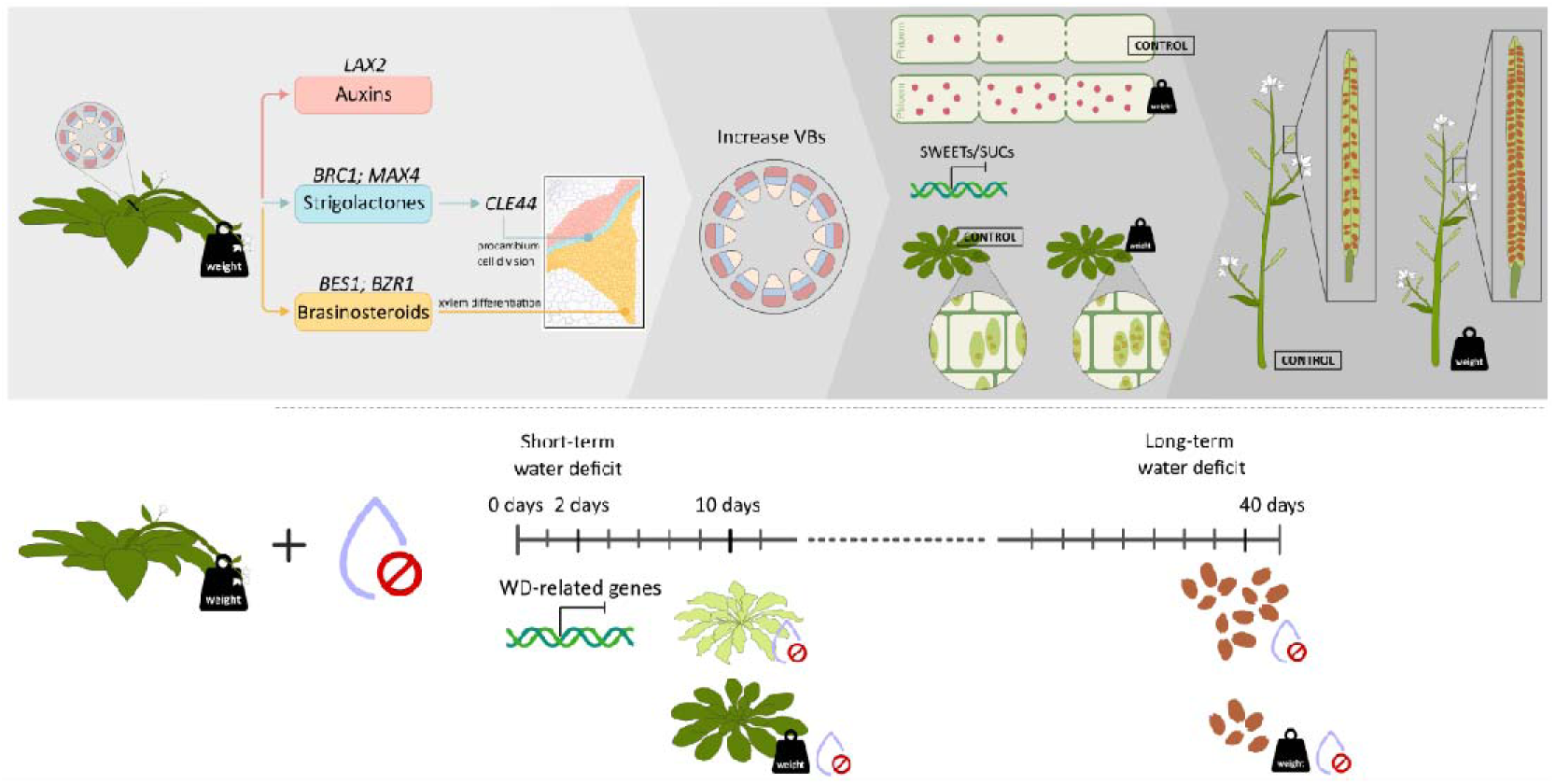
Regulatory network driving weight-induced vascular remodeling and yield enhancement. Weight treatment activates auxin transport (LAX2), brassinosteroid signaling (BES1/BZR1), and procambial regulators (BRC1, CLE44), promoting vascular bundle formation and xylem differentiation. Yield enhancement depends on functional sucrose transporters (SUC2, SWEET11/12/16) and dynamic regulation of starch metabolism, supporting efficient carbon redistribution to developing seeds. Mechanically treated plants display improved tolerance to short-term drought; however, prolonged water deficit during the reproductive stage reduces seed number, revealing a trade-off between enhanced vascular development and long-term hydraulic performance. Together, these coordinated pathways underlie the complex phenotype induced by weight treatment.

## EXPERIMENTAL PROCEDURES

### Plant material and growth conditions

*Arabidopsis thaliana* plants (Col-0 ecotype) were grown on Klasmann Substrat N° 1 compost (Klasmann-Deilmann GmbH, Germany) in a growth chamber at 22-24 °C under long-day (16/8hours light/dark cycles) conditions, with a light intensity of approximately 120 µmol m^-2^ s^-1^ in 8 × 7 cm pots. Four plants were planted per pot unless stated differently.

### Plant phenotyping

Plant architecture parameters were scored manually or with the aid of a ruler or gauge. All the experiments were performed with four to eight plants per genotype and per treatment, and repeated at least 3 times.

To determine the dry weight, fresh tissue was placed at 37 °C for four to six days until the weight was constant.

For determining seed yield, plants were sown one per pot, treated or not when they achieved 4.5 cm height, and then they grew normally until they were harvested at maturity when siliques were fully ripe. Yield was measured as seed weight (g) per plant.

### Histology and microscopy

Arabidopsis internodes were harvested and fixed as previously described (Cabello and Chan, 2019). Safranin-Fast green staining was used for xylem/phloem differential detection. After removing paraffin with 100 % xylene for 15 min at room temperature, sections were imbibed in ethanol (100%, 96%, and 90%), then the slices were transferred to Safranin (in ethanol 80%) for at least 4 hours. After that, slices were put in ethanol (90%, 96%, and 100%) to finally be imbibed on Fast-green dye (in ethanol 100%) for seconds and washed in ethanol 100% to remove excess dye. Slices were mounted on Canadian balsam.

For determining stem area, histological sections from at least four biological replicates were photographed and then measured using the ImageJ software (NIH, Maryland, United States).

### Soluble sugars, starch, and lipid content

For carbohydrate measurements, plant material (50-80 mg of each sample) was frozen and ground to powder in liquid nitrogen and extracted with a solution containing 5,4 mM phosphate buffer pH 7,5; 0,1 mM EDTA, 62,5% methanol, and 26,8% chloroform. The samples were incubated on ice for 20 minutes, and after that, 300 µl of water were added, mixed, and centrifuged for 5 min at 13,000 rpm. The supernatant was dried overnight at 40 °C and used to quantify sucrose and glucose. A fraction of this reaction was used to determine glucose content, and another fraction was treated with invertase to determine sucrose content. For starch extraction, the pellet was resuspended in 1 mL of ethanol and centrifuged at 13000 rpm for 5 minutes, dried at 70 °C, and finally resuspended in 250 µl of 0,1 N NaOH. Once a solution was obtained, 75 µl of 0,1 N AcH (pH 5,1) was added. Each sample (50 µl) was treated with amyloglucosidase overnight at 37 °C. Glucose was measured using a kit containing glucose-oxidase (GOD), peroxidase (POD), 4-aminofenazone (4-AF), phosphate buffer (pH 7), and 4-hydroxybenzene. The DO was measured at 505 nm, and glucose concentration was calculated using a glucose standard solution.

The extraction and quantification of lipids by gravimetry were based on the technique described by Siloto *et al*. (2006) and adapted for small volumes. Around 25 mg of seeds were ground and incubated with 400 µl of isopropanol for 10 min, at 65 °C, and then air-dried. Lipids were obtained after three extractions of a methanol, chloroform, and water biphasic solution (methanol: CHCl_3_:H_2_O). The first extraction was performed with 1.0 mL of methanol: CHCl_3_:H_2_O (2:2:1.8 [v/v]). The second and third extractions were performed with 1.0 mL of methanol:CHCl_3_:H_2_O (1:2:0.8 [v/v]). These three extractions were separated by centrifugations at 6700 g for 5 min. The lipid fractions were collected, and the solvents were completely evaporated. All the fractions were collected in a unique previously weighed microtube (W1), and after drying the sample, it was weighed again (W2). Lipids were expressed as W2 − W1/mg tissue. These experiments were repeated at least twice. Samples were obtained by pooling tissue from four to eight individual plants per genotype.

### RNA isolation and expression analyses by RT-qPCR

Total RNA for real-time RT-PCR was isolated from rosette leaves of 25-day-old plants using Trizol® reagent (Invitrogen, Carlsbad, CA, USA) according to the manufacturer’s instructions. Total RNA (1 μg) was reverse-transcribed using oligo(dT)_18_, and M-MLV reverse transcriptase II (Promega, Fitchburg, WI, USA).

Quantitative real-time PCR (qPCR) was performed using an Mx3000P Multiplex qPCR system (Stratagene, La Jolla, CA, USA) as described before (Cabello *et al*., 2016), using the primers listed in Supplementary Table ST3. Transcript levels were normalized by applying the ΔΔCt method. Actin transcripts (*ACTIN2* and *ACTIN8*) were used as internal standards. Four biological replicates, obtained by pooling tissue from 4 individual plants and tested in duplicate, were used to calculate the standard deviation. Each experiment was repeated at least three times.

#### RNA-Seq analysis

For RNA-seq analysis, total RNA was extracted from the stems of 4.5-cm tall Arabidopsis plants grown in soil, either untreated controls (C) or plants subjected to 6hours of weight treatment (T). Three biological replicates per genotype were sequenced by Novogene (Sacramento, CA, USA; Project number: H202SC24055447). Libraries were sequenced on the Illumina NovaSeq X-Plus platform, generating 150-bp paired-end reads with a minimum depth of 12 million reads per sample. RNA-Seq reads were analyzed on the Galaxy platform (Abueg et al., 2024). Briefly, adapter sequences were removed from the reads using Trimmomatic (Galaxy Version 0.39+galaxy2). The quality-filtered reads were aligned to the *Arabidopsis thaliana* reference genome (TAIR10) using RNA STAR (Galaxy Version 2.7.11a+galaxy1), employing a maximum intron length of 10,000 bp and guided by the gene and exon annotation from Araport V11. The read counts on each gene were then calculated using featureCounts (Galaxy Version 2.0.3+galaxy2). Differential expression analysis was performed using edgeR (Galaxy Version 3.36.0+galaxy5). Bigwig coverage files were generated using bamCoverage (Galaxy Version 3.5.4+galaxy0), and subsequently used for plotting with pyGenomeTracks (Galaxy Version 3.8+galaxy2). Functional enrichment analysis was performed using agriGO v2.0 (Tian et al., 2017). UpSet plots were generated to visualize intersections among gene sets responsive to weight (Ko et al., 2004 and this study), hypergravity (Tamaoki et al., 2009), or touch (Ku et al., 2019), using the UpSet diagram tool (Galaxy Version 0.6.5+galaxy2).

### CFDA Application

CFDA (6 mg/mL) was prepared in acetone and kept at −80°C. A 1:20 dilution in sterile water was used for application. The base of the inflorescences or flowers was immersed in CFDA until penetrated. All samples were excited by a 488 nm laser. Fluorescence 500 to 566 nm was monitored. Inflorescences and flowers were photographed after 5 or 10 minutes of application using an adapted Nikon camera. Quantification of CFDA transport was performed using ImageJ software.

### Statistical analyses

The data shown in Figures 1 to 6 were analyzed using a Student’s t-test or a two-way ANOVA. Student’s t-test was applied when comparing the expression of a specific gene or treatment, while in cases in which gene expression or phenotype was assessed across two treatments and more than one genotype, a two-way ANOVA was used. The data shown in the Figures represent a total of three replicates used to calculate the SE. For Student’s t-tests, differences were considered significant and indicated with asterisks when P-values were <0.05. For two-way ANOVA, when interaction terms were significant (P < 0.05), differences between means were analyzed using Tukey comparison and indicated by different letters.

## AUTHORS’ CONTRIBUTIONS

Conceived and designed the experiments: JVC and RLC. Performed most experiments: LR. Helped with some experiments: FAV. Analyzed the data: LR, MC, JVC and RLC. Wrote the paper: JVC, MC, and RLC.

## SUPPORTING INFORMATION

**Supplementary Figure S1. Weight-treated plants exhibit a slight yield penalty when grown at 60% field capacity**

**Supplementary Figure S2. Analysis of the overlap of genes upregulated or downregulated under different mechanical stress conditions**

**Supplementary Table S1. Lists of differentially expressed genes following weight treatment.**

**Supplementary Table ST2. Supplementary Table S2. Lists of enriched GO terms from differentially expressed genes in weight-treated plants.**

**Supplementary Table ST3. Oligonucleotides used for real-time qPCR.**

