## Supplementary figures and images for "Mechanical stress modulates source-to-sink partitioning and drought response in Arabidopsis"

### Supplementary Figure SF1

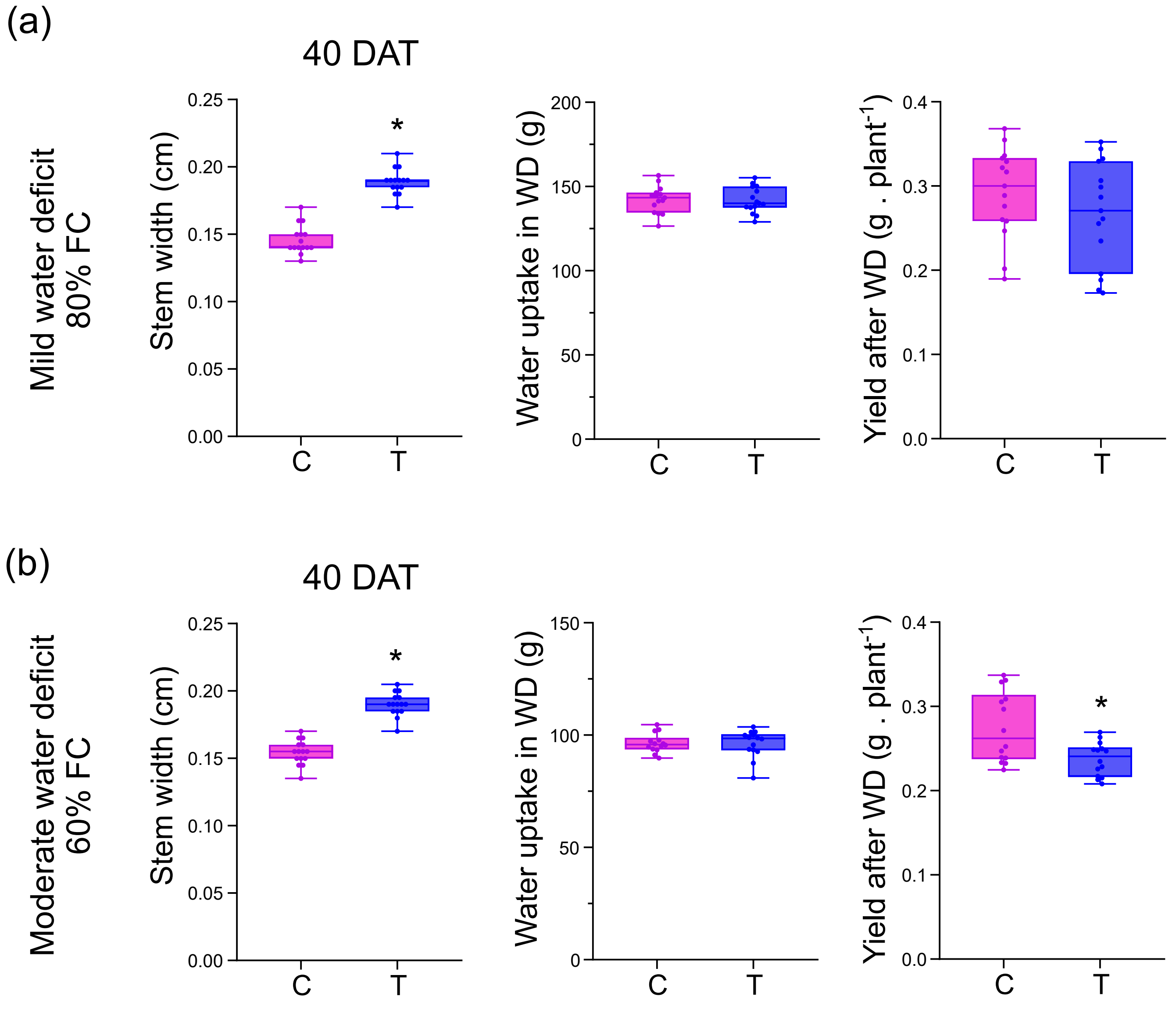

### Supplementary Figure SF2

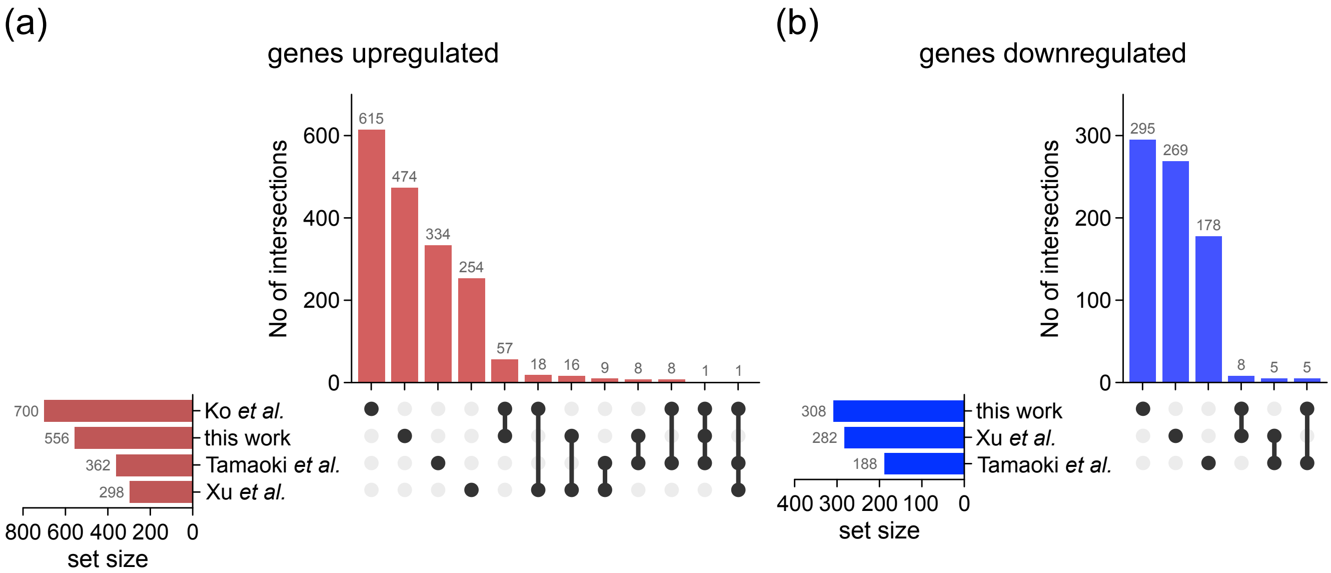
