## Supplementary Table ST3 for "Mechanical stress modulates source-to-sink partitioning and drought response in Arabidopsis"

**Supporting Figures**

**Supplemantary Table X:**

| Name | Function | Sequence 5’-3’ |
| --- | --- | --- |
| SWEET9-F | RT-qPCR | ACGCCGTCATGTGGTTCTTT |
| SWEET9-R | RT-qPCR | TGCTAGTTGGTTCTCTGTTGGC |
| SWEET10-F | RT-qPCR | AACTCCTTTGCCTTTGTCGTACA |
| SWEET10-R | RT-qPCR | GTCAACACGAAGATTGCGCC |
| SWEET11-F | RT-qPCR | AACAAGTGTACCTGCGGAAATGAT |
| SWEET11-R | RT-qPCR | AAGAGGACTGCTTGCCATGTTTAG |
| SWEET12-F | RT-qPCR | CATATGGCTCCTTTATGGTCTTGC |
| SWEET12-R | RT-qPCR | ACGTTTGGGAAGGCAACATAGATA |
| SWEET13-F | RT-qPCR | CTTCTACGTTGCCCTTCCAAATG |
| SWEET13-R | RT-qPCR | CTTTGTTTCTGGACATCCTTGTTGA |
| SWEET14-F | RT-qPCR | AAACGCTGTGGGATGCTTCA |
| SWEET14-R | RT-qPCR | TTCAAGAGCCCAAGAACCTTCA |
| SWEET15-F | RT-qPCR | CAATGACATATGCATAGCGATTCCAA |
| SWEET15-R | RT-qPCR | GGACTCATCACGACAATACTCTTAAG |
| SWEET16-F | RT-qPCR | GAGATGCAAACTCGCGTTCTAGT |
| SWEET16-R | RT-qPCR | GCACACTTCTCGTCGTCACA |
| SWEET17-F | RT-qPCR | AGTGACAACAAAGAGCGTGAAATAC |
| SWEET17-R | RT-qPCR | ACTTAAACCGTTGCTTAAACCAACC |
| SUC1-F | RT-qPCR | GACCTTTCGACGCCTTGTTC |
| SUC1-R | RT-qPCR | AATACTCCACTAATCGCCGCTG |
| SUC2-F | RT-qPCR | CATTCCTTTTGCACTAGCTTCCAT |
| SUC2-R | RT-qPCR | AGAACACCTAGGGAAAGTCCTTGG |
| SUC3-F | RT-qPCR | CAAGAACCGCAGCCGTAATC |
| SUC3-R | RT-qPCR | CTTGACCGCCACCGGAAT |
| SUC4-F | RT-qPCR | AGTGTCAAGCGAGGAACGCATA |
| SUC4-R | RT-qPCR | AGTCACACGAGAAGCCATTGC |
| PGM-F | RT-qPCR | GTGAAAGAGTATTGGGCGACA |
| PGM-R | RT-qPCR | CCGTGAACACAAACCGAACA |
| ADG1-F | RT-qPCR | CACCGTCTAAGATGCTTGATGC |
| ADG1-R | RT-qPCR | GATGTGCGAGTTTTTCCCAAT |
| ADG2-F | RT-qPCR | ATCAAGGAGAAACCTGCCACCA |
| ADG2-R | RT-qPCR | TCGTAGTAATCTGCCCCAAGC |
| GBSS-F | RT-qPCR | ATAGGGAGATTGGAGGAGCAGA |
| GBSS-R | RT-qPCR | CAATGTGGAAACCTGTGTAGC |
| SS1-F | RT-qPCR | CTTGATTACCAGAAGGGCATTG |
| SS1-R | RT-qPCR | CGTTTTCCCAAGAGTAGTTTCG |
| SS2-F | RT-qPCR | AGATAAAGCACGGGGATGGG |
| SS2-R | RT-qPCR | CCAACCAAGACCCGTTTCAC |
| SS3-F | RT-qPCR | CTGGGGCTGACTTTATTCTTGT |
| SS3-R | RT-qPCR | AGTCTTGCTCCATCACCGTCT |
| SS4-F | RT-qPCR | ACACGCCCTTAGAAAGCAGC |
| SS4-R | RT-qPCR | ACAAATCGGAGGCTGCGTAA |
| GWD-F | RT-qPCR | AACGAGAGAGCATACTTCAGC |
| GWD-R | RT-qPCR | CAATCGGTTTGCTTGGGTAG |
| AMY3-F | RT-qPCR | ATTATCATTCCGAGATTGCTGC |
| AMY3-R | RT-qPCR | CGGCTACAGACCAGTTTTGC |
| BAM1-F | RT-qPCR | AACGAGAGAGCATACTTCAGC |
| BAM1-R | RT-qPCR | CAATCGGTTTGCTTGGGTAG |
| BAM3-F | RT-qPCR | AACGAGAGAGCATACTTCAGC |
| BAM3-R | RT-qPCR | CAATCGGTTTGCTTGGGTAG |
| Actinas-F | RT-qPCR | GGTAACATTGTGCTCAGTGGTGG |
| Actinas-R | RT-qPCR | AACGACCTTAATCTTCATGCTGC |
| FLS1-F | RT-qPCR | GATTCGAAAGACATTGAAGGATACG |
| FLS1-R | RT-qPCR | CTCCGATAGCTTCTTCACATGCAC |
| RD28-F | RT-qPCR | CAACGTTTAAACTTAGCCACGA |
| RD28-R | RT-qPCR | CGTTGCAAAGATGAATTGAAAA |
| ERF53-F | RT-qPCR | AGCCAACTCCCGTGCATCAG |
| ERF53-R | RT-qPCR | CTACTGCCAATGCCGTGC |
| MAPKKK20-F | RT-qPCR | GATTGGGATTCTTTTACATTGG |
| MAPKKK20-R | RT-qPCR | ACCGGACTGTAAGCCAAC |
| PIP2B-F | RT-qPCR | CGCTAGAGACTCCCACGTTC |
| PIP2B-R | RT-qPCR | GAAACTCCTAGCCGGGTTG |
| NCED2-F | RT-qPCR | CGGTAACGGCGAAGAAAATG |
| NCED2-R | RT-qPCR | TGCCATGAAACCCATACGG |
| NCED5-F | RT-qPCR | GCTATTTGCGTTGAGCTACG |
| NCED5-R | RT-qPCR | TCGGAGAGCTTAAACACAACTT |
| ERD-F | RT-qPCR | CTCGTCTTCCGTCATCAGATC |
| ERD-R | RT-qPCR | CCCAACAAAAGGATTTGTGAGATG |
| Deh-F | RT-qPCR | GCCCAAGGAAGAGGAGAAGC |
| Deh-R | RT-qPCR | CCTCTGTTTCACATTGATCTTCAGC |
| NAC019-F | RT-qPCR | TCAGCAACAACGGTACTTCG |
| NAC019-R | RT-qPCR | TGCGGTTTGGGTTAGAAAAC |
